# Predation limits the abundance of Δ*gacA* cheaters in *Pseudomonas protegens* populations

**DOI:** 10.1101/2025.11.08.687362

**Authors:** Margaret I. Steele, Emily E. Oxender, Carol Ge, David C. Queller, Joan E. Strassmann

## Abstract

In bacteria, defense against predators and competitors often requires cooperation within large clonal bacterial populations. Mutations that inactivate biosynthesis of costly molecules involved in cooperative defense can increase the growth rate of the mutants, while mutant cells continue to benefit from the efforts of wild type cells. The GacA-GacS signaling pathway regulates biosynthesis and secretion of toxic secondary metabolites and proteins that play crucial roles in bacterial defense in *Pseudomonas* species, including resistance to the protozoan predator *Dictyostelium discoideum*. A *P. protegens* Δ*gacA* mutant is vulnerable to predation but can benefit from protection provided by wild type cells in mixed populations. Wild type *P. protegens* sporadically infects *D. discoideum* fruiting bodies which may aid in bacterial dispersal to new environments. We found that the Δ*gacA* mutant does not infect fruiting bodies and that the wild type does not infect when at low abundance in the population. The *P. protegens ΔgacA* mutant is a cheater whose fitness under predation depends on its relative abundance. Protozoan predators can limit the abundance of Δ*gacA* mutants in *Pseudomonas* populations, preventing selective sweeps that could lead to the loss of the important but costly Gac regulon.

## Introduction

Bacteria in soil ecosystems face predation from protists and nematodes, as well as competition from fungi and other bacteria. To protect themselves, many bacteria secrete toxic proteins and metabolites (1–4). Bacterial defense mechanisms are often cooperative in nature and depend on millions of cells working together to produce compounds that benefit the population as a whole. Though some defensive compounds are produced through interspecific cooperation (5), production of public goods is most effective as an evolutionary strategy when the populations are clonal or nearly so. Biosynthesis of defensive molecules can be costly and individuals with mutations that inactivate production of these compounds may have higher growth rates, allowing them to outcompete wild type cells while still receiving the protection provided by the wild type population. These mutants are cheaters, which, by definition, exploit public goods produced by closely related individuals and, by not contributing to the effort, reduce wild type fitness.

Overabundance of cheaters can reduce WT fitness and eventually eliminate cooperative behaviors. For example, in biofilms of *Bacillus subtilis*, cheaters can arise that do not produce energetically costly exopolysaccharides that help make up the biofilm matrix. In a short-term response to cheaters, matrix producers can increase levels of matrix expression, but exopolysaccharide-deficient cheaters tend to take over in mixed populations and abolish biofilm formation (6). In *Myxococcus xanthus*, mutants deficient in extracellular protein production—essential for multicellular development—cheat on the wild type by forming spores at greater rates. Short range signaling limits cheating in wild type and mutant mixtures by requiring sufficient wild type cells (7). Cheaters often experience the highest fitness when they are present at low abundance (frequency-dependent selection), as is the case for *Pseudomonas aeruginosa* cheaters that do not produce siderophores that are needed to scavenge iron from the environment (8). External factors can limit the spread of cheaters within a population. Proliferation of *Pseudomonas protegens* Gac-mutants in liquid culture can be constrained by co-culture with a bacterial competitor (9). Another study on *P. aeruginosa* found that adding the toxic phenazine pyocyanin to liquid cultures limited the spontaneous proliferation of protease-deficient cheaters (Castañeda-Tamez et al., 2018).

The protist *Dictyostelium discoideum* preys on diverse bacteria and sometimes serves as a host for bacterial pathogens. During the vegetative stage of *D. discoideum*’s life cycle, unicellular amoebae use chemotaxis to hunt bacteria which they then consume through phagocytosis (10). When prey availability is low, the onset of starvation triggers amoebae to aggregate and form a multicellular motile slug. The slug develops into a fruiting body that aids in the dispersal of spores by insects (11). Bacteria employ numerous strategies to defend themselves against predation by *D. discoideum* and other protists. Many predation-resistant bacteria have been isolated from wild *D. discoideum* populations, including several *Pseudomonas* species (12, 13). Some of these unpalatable *Pseudomonas* infect *D. discoideum* fruiting bodies (12, 14), which may aid in dispersal of bacteria to new environments. Though potential defense genes, such as protein secretion systems and secondary metabolite biosynthetic pathways, vary between species (14), the GacS-GacA two-component system that regulates many of these genes is required for predation resistance in diverse *Pseudomonas* lineages (2, 15–18).

The GacA-GacS two-component system regulates hundreds of genes in *Pseudomonas*, including many that are likely to contribute to predation resistance, such as synthesis of secondary metabolites and exoprotease, secretion systems, and biofilm production (19). The GacS sensor histidine kinase responds to an environmental signal by phosphorylating the response regulator GacA. GacA activates transcription of non-coding small RNAs, which bind to RsmA to derepress translation of genes in the regulon (20). Gac^−^ mutants are often detected in both laboratory and natural *P. protegens* populations. Whether these mutants act as cheaters appears to depend on the context of the interaction. *P. protegens ΔgacA* mutants are non-motile cheaters that only swarm when mixed with wild type (21). In contrast, Δ*gacA* mutants, which exhibit reduced biofilm production in monoculture, increase biofilm formation by wild type in mixed populations, possibly by supplying extra siderophores (22). The presence of a bacterial competitor reduces the detection of spontaneous *P. protegens* Gac^−^ mutants in laboratory culture (9), suggesting that biotic interactions can constrain the spread of such mutants. A previous study isolated nearly identical GacA^+^ and GacA^−^ *P. protegens* (previously identified as *Pseudomonas fluorescens*) strains from a single naturally infected *D. discoideum* strain (16), leading us to hypothesize that GacA mutants may also act as cheaters in the context of predation resistance by benefiting from defensive compounds produced by wild type bacteria.

In this study, we investigated how predation by *D. discoideum* affects competition between wild type and *ΔgacA P. protegens*. To determine whether *P. protegens ΔgacA* acts as a cheater, we measured the effect of mutant abundance on wild type and Δ*gacA* fitness with and without predation. Finally, we investigated the effect of initial mutant frequency on the proportion of *D. discoideum* fruiting bodies that become infected with WT or *ΔgacA P. protegens*, since *D. discoideum*-facilitated dispersal may also contribute to bacterial fitness. We found that Δ*gacA* better survived predation when it was at a lower abundance in the population and that high mutant abundance compromised wild type dispersal but not resistance to predation. Furthermore, predation limits mutant abundance, and may allow WT and Gac^−^ strains to coexist despite the growth advantage observed in Gac^−^ strains.

## Results

### *P. protegens ΔgacA* acts as a cheater in the presence of predatory amoebae

We predicted that Gac^−^ mutants, which are susceptible to predation by D. discoideum amoebae, are protected from predation by WT production of Gac-regulated defensive compounds. To test this hypothesis, we cultured WT *P. protegens* and an isogenic Δ*gacA* deletion mutant at different starting frequencies with and without *D. discoideum*. We hypothesized that the mutant would have higher relative fitness when at low frequency in the presence of predatory amoebae. If predation resistance is dependent on cooperative secretion of defensive compounds such as secondary metabolites or exoprotease, we expected to see a decrease in wild type relative fitness in the presence of amoebae when mutant abundance is high. Secondary metabolism imposes a metabolic cost on the cell by diverting energy that could otherwise be put into growth and replication and we observed that spontaneous Gac^−^ mutants rapidly increase in frequency in *P. protegens* liquid cultures. Due to this growth advantage, we predicted the Δ*gacA* mutant would outcompete the WT in the absence of predation.

To determine how predation affects mixed populations of WT and Δ*gacA* cells, we cultured the two strains at starting frequencies of 100:0, 99:1, 50:50, 1:99, and 0:100 on nutrient agar with and without *D. discoideum* amoebae, then quantified the number of each strain remaining after 7 days (Figure 1A). We log transformed the recovered CFU and used three-way linear mixed effects models to measure how the presence of amoebae contributed to the effect of each strain on the other. In the linear mixed effects model measuring the effect of WT on Δ*gacA*, we excluded the 0% Δ*gacA* control group and treated the starting frequency and the presence or absence of amoebae as fixed effects and the day on which the experiment was performed as a random effect (log Δ*gacA* CFU ~ starting frequency * amoebae + (1 | Experiment). To measure the effect of ΔgacA on WT, we used a similar linear mixed effects model and excluded the 0% WT control group. Since the relationship between starting ratio and log transformed recovered CFU is non-linear, we treated the starting ratio as a factor instead of numeric.

**Figure 1.**
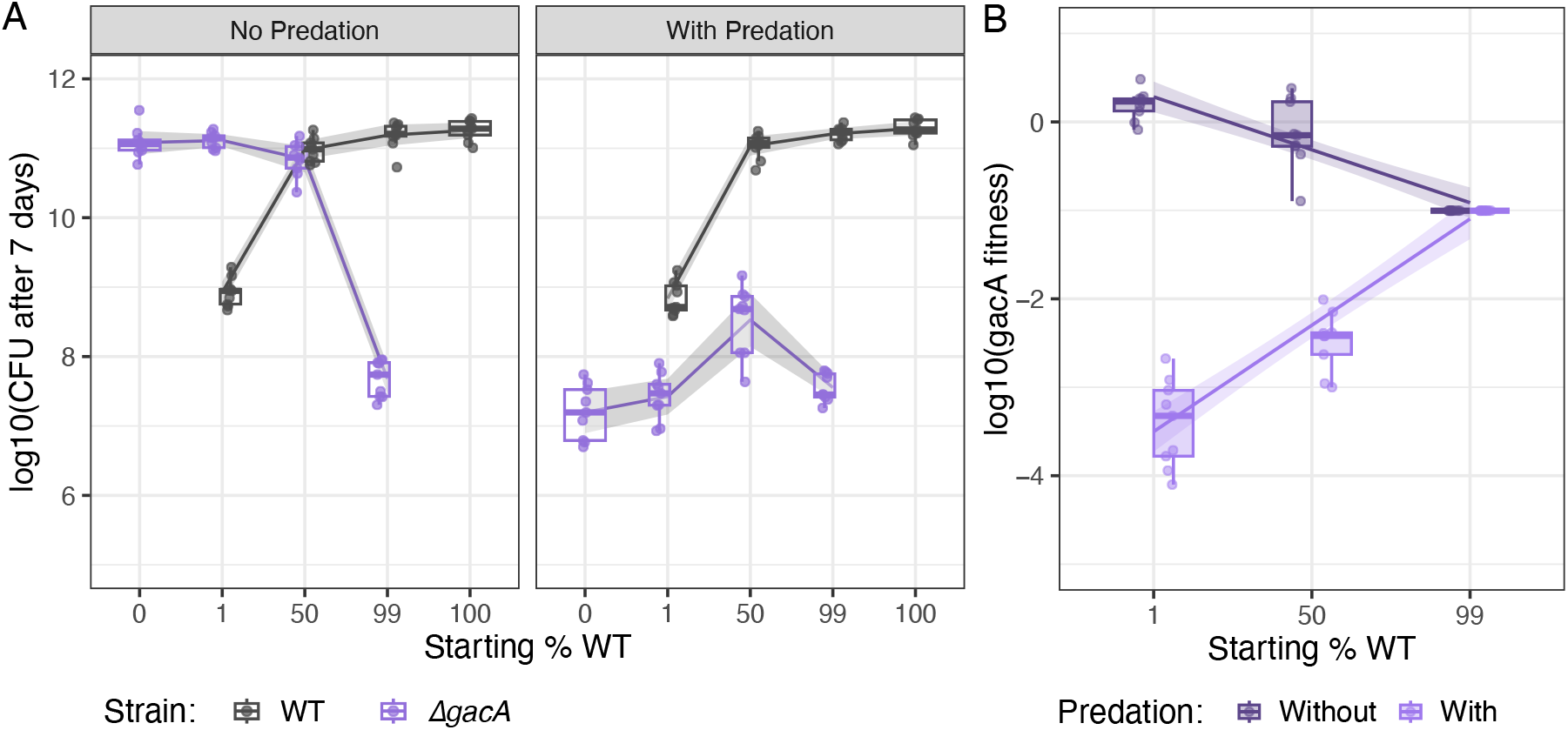
Predation by *D. discoideum* dramatically reduces the prevalence of *P. protegens* Δ*gacA*, but this effect decreases as the frequency of WT increases. (A) WT and Δ*gacA* CFU remaining after 7-day coculture with and without *D. discoideum* QS157. Points represent CFU counts from replicate plates (3 independent experiments, 3 technical replicates per experiment). (B) Relative fitness of the Δ*gacA* mutant at different starting frequencies with and without predation. Relative fitness was calculated as ((final frequency/(1 – final frequency))/(starting frequency/ (1 – starting frequency)).

The presence of amoebae greatly reduces the number of Δ*gacA* CFU recovered (F_1,64_=1216.5, p < 2×10^−16^) and the effect of the starting frequency on final Δ*gacA* CFU depends on whether amoebae are present (F_3,64_=145.8, p < 2×10^−16^), with higher concentrations of WT reducing the effect of predation on Δ*gacA* (Figure 1A). There was no overall effect of amoebae on the number of WT CFU recovered and the presence of amoebae did not affect how WT performs at different starting ratios. When *D. discoideum* was present, we obtained the highest Δ*gacA* yields when the WT and mutant started at equal frequency, balancing the negative effects of competition against the benefit of protection. WT yields are essentially identical in control and *D. discoideum* treatments. In the absence of predation, the initial frequency of WT has a slight negative effect on the relative fitness of Δ*gacA* (estimate = −0.012±0.0015 SE), reflecting competition with WT (Figure 1B). When amoebae are present, there is a positive effect of starting abundance of WT on Δ*gacA* relative fitness (estimate = 0.037±0.0022 SE), with fitness improving with WT frequency. However, Δ*gacA* relative fitness is always less than 0 in the presence of amoebae, showing that it is at a competitive disadvantage to WT. At a starting ratio of 99% WT, the Δ*gacA* mutant is essentially unaffected by predation. This is likely because there are insufficient prey bacteria to support *D. discoideum* growth and so the abundance of the predator is low.

### The presence of additional prey bacteria does not alter the overall trend

The outcomes of predator-prey interactions between *D. discoideum* and bacteria are often determined by the ratio of predators to prey. Some bacteria are susceptible to predation at low density but resistant to predation at high density (23, 24). *D. discoideum* cannot grow and does not form fruiting bodies if the ratio of predation resistant to prey bacteria is too high (14), so in treatments with 100% or 99% WT *P. protegens*, Δ*gacA* cells likely experience reduced predation pressure. To examine how predation effects interactions between WT and Δ*gacA* with similar numbers of amoebae in each treatment, we supplemented the cultures with the prey bacterium *Klebsiella pneumoniae*. Each culture was inoculated with the same total number of bacteria, but the mixture contained 90% *K. pneumoniae* and 10% *P. protegens* (WT + Δ*gacA*).

When the prey bacteria *K. pneumoniae* was present, the presence of amoebae strongly affected the number of Δ*gacA* CFU recovered (F_1,120_=3195.93, p<2.2×10^−16^) and the effect of WT frequency on Δ*gacA* depended on whether amoebae were present (F_5,120_=3.264, p=0.0085) (Figure 2A). Surprisingly, the presence of amoebae did reduce the number of WT CFU recovered (F_1,119_=13.55, p=0.00035) and altered the effect of Δ*gacA* on WT (F_5,119_=6.32, p=3.076×10^−5^). Only when WT starts at very low frequency (0.1%) does it do better in the treatment with amoebae than in the control (estimate = −0.270±0.799 SE, t_119_=-3.38, p = 0.001). When we looked at relative fitness of each *P. protegens* strain in the presence of *K. pneumoniae*, the presence of amoebae strongly affected Δ*gacA* fitness (F_1,109_=1490.43, p < 2.2 x 10^−16^) and the effect of WT on Δ*gacA* fitness depends on whether amoebae are present (F_1,109_ = 241.33, p<2.2 x 10^−16^) (Figure 2B). There is a small but significant negative effect of starting frequency of WT on Δ*gacA* relative fitness in the absence of amoebae (estimate = −0.069 ± 0.012 SE) and a positive effect of WT frequency on Δ*gacA* relative fitness in the presence of amoebae (estimate = 0.250 ±0.016 SE). Since *K. pneumoniae* is present as an alternative food source, Δ*gacA* relative fitness is significantly lower in the presence of amoebae, even with a high starting frequency of WT.

**Figure 2.**
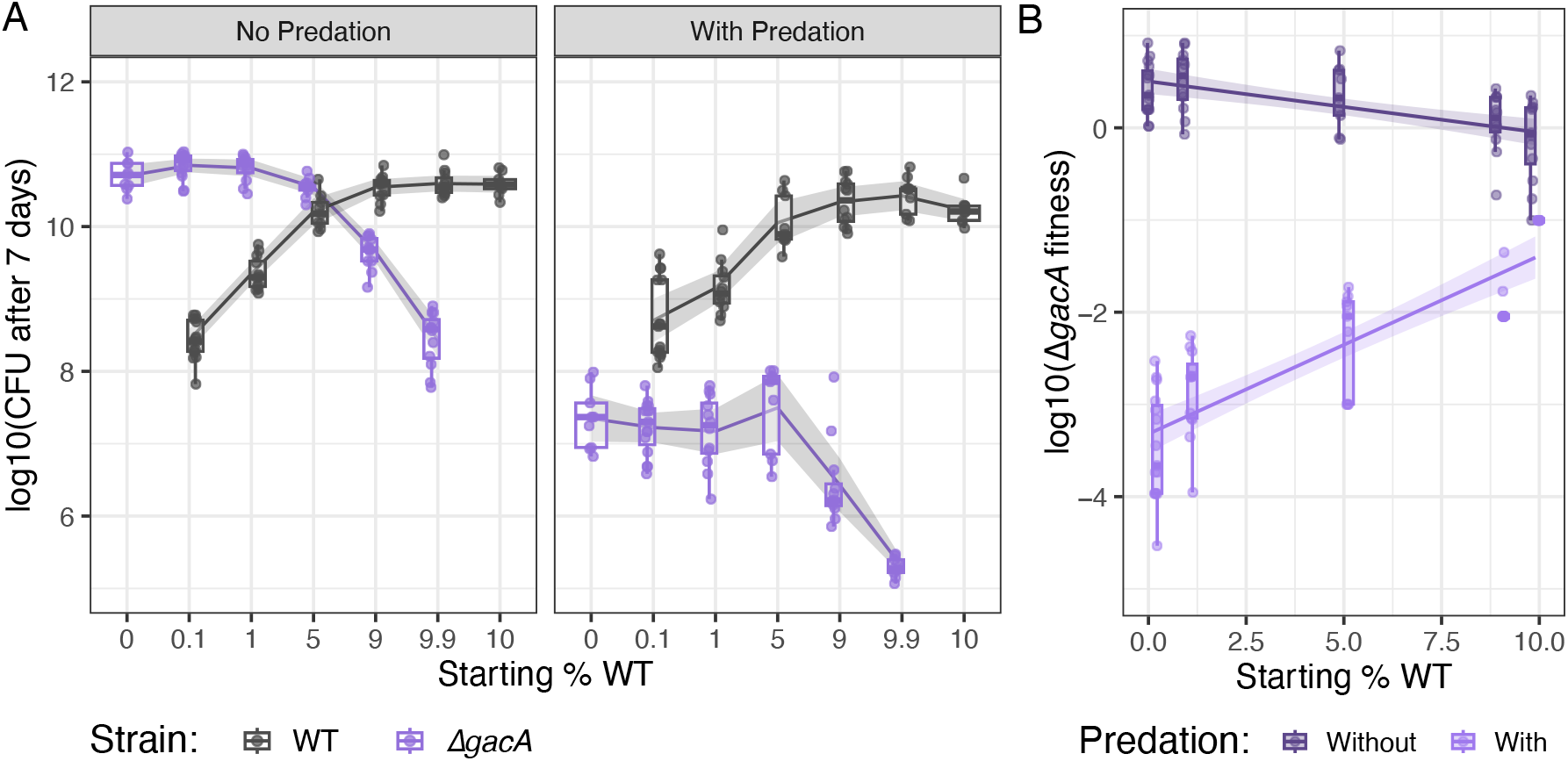
When additional prey bacteria are available, predation by *D. discoideum* reduces the prevalence of Δ*gacA*, but this effect is weakened as the frequency of WT increases. (A) WT and Δ*gacA* CFU remaining after 7-day coculture with and without *D. discoideum*. Points represent CFU counts from replicate plates (3-4 independent experiments with 3 technical replicates per experiment). (B) Relative fitness of the Δ*gacA* mutant at different starting frequencies with and without predation.

### Coinfections of *D. discoideum* by wild type and Δ*gacA P. protegens* are rare

*P. protegens* WT sporadically infects *D. discoideum* fruiting bodies (12, 14, 16). A previous study reported that a *P. protegens* strain with a nonsense mutation in *gacA* and a nearly identical strain with a functional *gacA* gene were co-isolated from a naturally infected *D. discoideum* fruiting body (16). Based on this, we hypothesized that the Δ*gacA* knockout may retain the ability to infect *D. discoideum* fruiting bodies despite being susceptible to predation or may cooperatively infect when WT is present. Contrary our expectations, the Δ*gacA* mutant was rarely detected in fruiting bodies, even when the WT was present (Figure 3). In all experiments, we detected WT P. *protegens* in more infected fruiting bodies when WT started at a higher frequency in the population. A binomial logistic mixed model revealed a significant interaction between strain and starting frequency (beta = 0.62±12, z = 5.412, p = 6.22e-08), indicating that the fraction of fruiting bodies infected by WT increased with the initial frequency of WT in the population, but no such relationship was detected for Δ*gacA*. As expected, no bacteria were detected in fruiting bodies grown on 100% *K. pneumoniae*.

**Figure 3.**
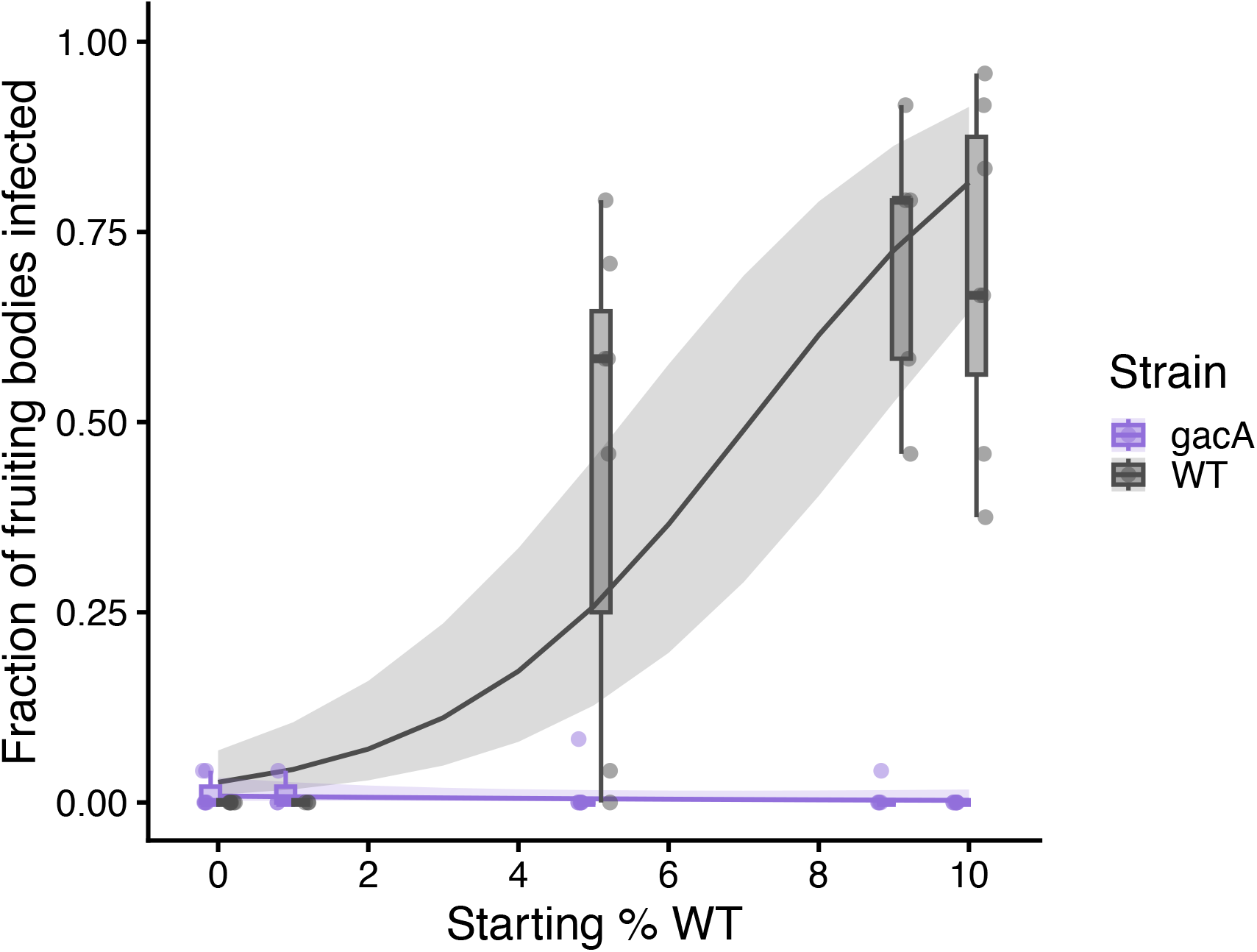
WT *P. protegens* infects a higher proportion of *D. discoideum* fruiting bodies when it starts at a higher frequency in the population, while Δ*gacA* rarely infects at any frequency. Each point represents the fraction of *D. discoideum* fruiting bodies (of 24 sampled) in which P. *protegens* was detected in an independent experiment. Box plots show the median and range of the data. The lines represent the fraction of infected fruiting bodies based on a logistic mixed-effects model with a 95% confidence interval.

## Discussion

*Pseudomonas* species engage in a variety of cooperative behaviors that allow them to survive biotic and abiotic stressors. Many of these behaviors are regulated by the GacS-GacA two-component system, which regulates up to 10% of all annotated genes in *Pseudomonas fluorescens* Pf-5 (19). GacS and GacA mutants are found in natural *Pseudomonas* populations (16, 25, 26) and recent work has shown spontaneous mutations in the *gacA* and *gacS* genes allow *P. protegens* Pf-5 to rapidly outcompete WT when grown in rich medium in the lab (9). These strains are deficient in biofilm formation (22), swarming motility (21), production of secondary metabolites (9) and production of exoprotease. In *P. protegens* Pf2, both iron limitation and predation by the soil amoeba *D. discoideum* affect competition between WT and an isogenic Δ*gacA* knockout (27). *P. protegens* CHA0 *gacS* mutants are preferentially consumed and show negative-frequency dependent selection in the presence of predatory nematodes and *Acathamoeba castellanii* amoebae (28). Gac^−^ mutants have been shown to act as cheaters during swarming (21) but not biofilm formation (22). However, whether Gac^−^ strains act as cheaters in the context of predation resistance has not been established.

*P. protegens* Gac^−^ mutants have a competitive advantage over WT in liquid culture (9) but suffer increased susceptibility to protozoan grazers (2, 15, 17). We used the bactivorous amoeba *D. discoideum* to investigate whether *P. protegens* Δ*gacA*, which has a Gac^−^ phenotype, acts as a cheater by exploiting Gac-regulated defensive compounds produced by WT. We tested the effects of increasing Δ*gacA* mutant abundance on both mutant and WT fitness in the presence of the predator. We predicted that Δ*gacA* would outcompete WT in the absence of predation due to increased growth rate, that Δ*gacA* would be more susceptible to predation when it was at high frequency in the population, and that high frequencies of Δ*gacA* would reduce WT fitness because resistance is partially determined by population size (23, 24). We considered two different effects on WT fitness: susceptibility to predation and loss of *D. discoideum*-facilitated dispersal (14). We performed fitness experiments with and without the addition of the prey bacterium *Klebsiella pneumoniae*. The inclusion of *K. pneumoniae* ensures *D. discoideum* amoebae are growing even when the *P. protegens* population is dominated by the inedible WT but also introduces interspecific competition. However, preferential feeding on *K. pneumoniae* (29) may help to minimize the effects of interspecific competition.

We found that the frequency of WT strongly affected Δ*gacA* survival in the presence of amoebae, but Δ*gacA* did not outcompete WT in the absence of predation or make WT more susceptible to predation when at low frequencies. The positive correlation between Δ*gacA* fitness and WT frequency suggests that the mutant benefits from WT defense mechanisms. The inverse correlation between Δ*gacA* starting frequency and relative fitness is very similar to the negative frequency-dependent selection seen in *P. protegens* CHA0 *gacS* mutants subject to predation by nematodes or protists (28). Though Δ*gacA* had the highest relative fitness when it started at low frequency, it yielded the highest CFU when it started at a frequency of 50%, suggesting a balance between predation and intraspecific competition. Interestingly, this trend holds true regardless of whether additional prey bacteria are present. When WT abundance is high and no *K. pneumoniae* is present, Δ*gacA* has the same relative fitness with or without the addition of amoebae, likely because the high concentration of WT *P. protegens* and the lack of available prey bacteria inhibits *D. discoideum* growth, resulting in little to no predation. However, Δ*gacA* relative fitness is always less than 0 when amoebae are present, indicating a competitive disadvantage against WT. Surprisingly, even when starting at a high initial frequency with no amoebae, Δ*gacA* relative fitness is not much more than 0, suggesting that it does not have a competitive advantage over WT. This contradicts the results of previous stationary phase liquid culture experiments (9), but it is possible increased population structure in cultures grown on agar reduce the effects of the growth advantage.

High concentrations of cheaters can dramatically affect the fitness of WT bacteria in the presence of predators. For example, predatory nematodes will consume WT *P. fluorescens –* which depends on the production of chemorepellents to evade predation – if it is mixed with a *gacS* mutant at a ratio equal to or less than 1:5 (1). Over time, increased prevalence of cheater mutations can cause species that rely on cooperative behaviors to resist predation to adopt less effective individualistic defense strategies, as is seen in *Pseudomonas putida* populations co-evolved with a predatory flagellate (30). WT *P. protegens* was unaffected by *D. discoideum* in our *Pseudomonas*-only experiment, even when it started at low frequency. However, when *K. pneumoniae* was added we observed small but significant reductions in the number of WT CFU recovered from treatment groups with amoebae. This could be caused by interspecific competition limiting *P. protegens* growth, but it is likely because *K. pneumoniae* is an excellent food source for *D. discoideum* and allows the predator population to reach higher densities (29). This is an example of an Allee effect, in which higher population size results in higher individual fitness. Our observation that WT *P. protegens* is not more vulnerable to predation when it starts at a low initial frequency somewhat contradicts previous studies that found that large population size was necessary for *Pseudomonas fluorescens* HKI0770 (24) and some *Escherichia coli* strains (23) to resist grazing by *D. discoideum*. These earlier studies restricted population size by limiting nutrient availability, while we limited population size through intraspecific competition. A more direct comparison of these two methods may be warranted to detangle the influences of nutrient depletion and Allee effects on bacterial resistance to predation.

One possible explanation for this discrepancy is the time it takes for spores to hatch into amoebae that can start consuming bacteria. When WT starts at low frequency, this initial period of bacterial growth without predation would introduce population structure with small patches of WT bacteria growing amidst a lawn of Δ*gacA* cells, which could exclude Δ*gacA* from protection by secreted compounds that do not diffuse far though agar. Even when WT *P. protegens* starts out at 0.1% of the population (approximately 2.4 x 10^5^ cells per plate), it may reach a high enough local density for cooperative defense mechanisms to be effective. Alternatively, WT *P. protegens* cells may utilize additional individual defenses that make them non-preferred prey even when they are in the minority. Regulatory pathways that control cooperative behaviors can also contribute to private defenses. For example. *Pseudomonas aeruginosa* mutants defective in quorum sensing are vulnerable to predation by *Tetrahymena pyriformis* and are unable to exploit resistance of wild type bacteria (31).

*D. discoideum* has a primitive immune system used to eliminate potentially pathogenic bacteria during its multicellular developmental cycle (32–34), which ensures that most fruiting bodies are free from bacteria that might affect the fitness of the spores. Despite this, some bacteria end up inside *D. discoideum* fruiting bodies (12, 13, 16). *P. protegens* is one of several *Pseudomonas* species that sporadically infect fruiting bodies (14). During multicellular development, *D. discoideum* cells aggregate to form a motile slug, which can travel several centimeters before developing into a fruiting body, which facilitates dispersal of spores by invertebrates (11). The ability to infect fruiting bodies may provide a fitness advantage for bacteria by allowing them to disperse away from closely related competitors. Other *Pseudomonas* species allow fruiting bodies to be infected by bacteria that are otherwise edible in addition to themselves, but this does not appear to be true for *P. protegens* (14). Although nearly identical Gac^+^ and Gac^−^ *P. protegens* strains were previously co-isolated from *D. discoideum* in the wild (16), we found no evidence that *P. protegens* Δ*gacA* can infect fruiting bodies in the lab with or without the assistance of WT *P. protegens*. Coinfection may be a very rare occurrence that persists by providing unique advantages. For example, Gac^+^ and Gac^−^ *Pseudomonas* strains cooperatively form biofilms in which the Gac^+^ cells produce phenazines and the Gac^−^ cells produce siderophores (22). We also observed that the WT does not infect fruiting bodies when its starting abundance is low. Therefore, carriage likely depends on population density. Furthermore, the discovery that WT can infect fruiting bodies while Δ*gacA* cannot suggests that *D. discoideum*-facilitated dispersal may be one way for *P. protegens* populations to escape cheaters. Further work is needed to better understand the effects of *D. discoideum*-facilitated dispersal on bacterial fitness.

This study provides new insight into how Gac^−^ mutants, which are frequently found in wild and laboratory *Pseudomonas* populations, stably coexist with WT in nature despite their apparent fitness advantage in the lab. *Pseudomonas* species are commonly used for biocontrol of plant pathogens due to their ability to produce a variety of antimicrobial secondary metabolites, which are positively regulated by GacS and GacA (2, 18, 35). Elimination of secondary metabolite production due to excessive proliferation of Gac^−^ cheaters is a problem for biocontrol efforts and would dramatically alter microbial ecosystems. Interactions with predators like *D. discoideum* likely help to prevent proliferation of Gac^−^ cells in nature.

## Methods

### Strains and culture conditions

Antibiotics were added to LB agar at the following concentrations: 20 μg/mL gentamicin (Gm) and 30 μg/mL kanamycin (Km). For each experiment, fresh cultures were started by streaking cells from glycerol stocks onto LB agar (LB broth, Miller (Fisher) and 15 g of agar (Fisher Scientific) per L) and incubating them at 30ºC. Bacterial dilutions were prepared using KK2 (2.25 g KH_2_PO_4_ (Sigma Aldrich) and 0.67 g K_2_HPO_4_ (Fisher Scientific) per L). For assays including *D. discoideum*, cells were plated on SM/5 agar (2 g glucose (Macron), 2 g bacteriological peptone (Oxoid), 2 g Bacto Yeast Extract (Gibco), 0.2 MgSO_4_•7H_2_O (Fisher Scientific), 1.9 g KH_2_PO_4_ (Sigma Aldrich), 1 g K_2_HPO_4_ (Fisher Scientific), and 15 g of agar (Fisher Scientific) per L). *D. discoideum* cultures were incubated at room temperature (20-22°C).

**Table 1:** Strains used in this study.

| Species | Strain | Description | Source |
| --- | --- | --- | --- |
| *Klebsiella pneumoniae* | Kp |  | (Dictybase stock center) |
|  | 18P_8.2_Bac1-GFP | Chromosomal insertion of GmR and GFP at attTn7 site | (Steele et al., 2023) |
| | 18P_8.2_Bac1ΔgacA | Replacement of *gacA* with KmR | (Steele et al., 2024) |
| *Dictyostelium discoideum* | QS157 | Wild clone | (Brock et al., 2011) |

### Fitness assays

Fitness assays were conducted to determine the effect of predation on mixtures of wild type and Δ*gacA* bacteria. *P. protegens* 18P_8.2_Bac1-Gm-GFP (wild type) and 18P_8.2_Bac1 ΔgacA were streaked out from glycerol stocks on LB agar and incubated at 30°C for 1 day. After one day, each bacterium was diluted in KK2 buffer to an OD_600_ of 1.5. For assays that included food bacteria *Klebsiella pneumoniae* (Kp), bacterial mixtures were made using 90% Kp, and varying ratios of wild type and mutant bacteria: 10% WT, 9.9% WT, 9% WT, 5% WT, 1% WT, 0.1% WT, and 0% WT. For assays without Kp, bacteria were mixed at the following ratios: 100% WT, 99% WT, 50% WT, 1% WT, and 0% WT. For the predation condition, *D. discoideum* QS157 spores were collected from fruiting bodies on SM/5 plates, counted using a hemocytometer, and then diluted to a concentration of 2 x 10^6^ spores/mL. For controls with bacteria alone, a 200 µl volume of each mixture was spread in triplicate on SM/5. For the condition with *D. discoideum* predation, a 200 µl volume of each bacterial mixture was mixed with 50 µl of *D. discoideum* spores. All plates were incubated at room temperature for 7 days. To quantify CFU for the starting bacterial mixtures, seven 1:10 serial dilutions were prepared and 10 µl droplets of each dilution were spotted in triplicate on LB supplemented with kanamycin and LB supplemented with gentamicin. The plates were incubated at 30°C, and CFU were counted after 1-2 days. After 7 days, plates were washed with 5 mL of KK2 and then scraped to dislodge cells. The same dilution process was performed with the recovered cells to quantify final CFU.

### Co-infection assay

Co-infection assays were performed to determine whether *P. protegens ΔgacA* is capable of infecting D. discoideum fruiting bodies alone or in the presence of WT. *P. protegens* 18P821-Gm-GFP (WT), 18P821ΔgacA::KmR, and *K. pneumoniae* were grown overnight on non-selective LB plates, collected into KK2, then diluted to an OD of 1.5. Cells were mixed to 90% *K. pneumoniae* with 10% WT, 9% WT, 5% WT, 1% WT, or 0% WT (with the remaining percentage made up by ΔgacA). A negative control was included in which *D. discoideum* was grown with *K. pneumoniae* alone. *D. discoideum* spores were collected, counted using a hemacytometer, and diluted to a concentration of 8×10^6^. 200 µl of bacterial mixtures and 50 µl of the spore solution were spread on replicate SM/5 agar plates. After 7 days, sterile 10 µl pipet tips were used to collect 24 individual fruiting bodies from each treatment group, which were then transferred to wells of a 96-well plate containing 30µl KK2. The plate was shaken gently to dislodge bacteria from fruiting bodies. 5 µl droplets from each well were spotted on LB Gm, LB Km, and non-selective LB to determine the prevalence of WT, Δ*gacA*, and *K. pneumoniae* infections.

### Statistics

Statistical analyses were performed in R (version). CFU counts from fitness assays (with and without *K. pneumoniae*) were log10-transformed and then analyzed using three-way linear mixed-effects models (lme4 and lmerTest packages) in which the starting frequency of WT and the presence or absence of amoebae were treated as fixed effects and the date of the experiment as a random effect. Separate analyses were performed for WT and Δ*gacA* CFU, excluding the 0% control group, to measure the effect the intraspecific interaction on the final abundance of each strain. The significance of fixed effects was estimated using a Type III ANOVA with Satterthwaite’s approximation for denominator degrees of freedom. Relative fitness of Δ*gacA* was calculated using the equation ((final frequency/(1 – final frequency))/(starting frequency/ (1 – starting frequency)). Linear mixed-effects models were used to measure the effects of initial WT frequency and amoeba presence with date as a random effect. Model significance and pairwise contrasts were assessed using estimated marginal means (emmeans package) with 95% confidence intervals. The proportion of *D. discoideum* fruiting bodies infected by WT or Δ*gacA P. protegens* was analyzed using a binomial logistic mixed-effects model, using the glmer() function in lme4.

